# Molecular understanding of anthocyanin biosynthesis activated by PAP1 in engineered red *Artemisia annua* cells and regulation of 2, 4-dichlorophenoxyacetic acid

**DOI:** 10.1101/2023.03.17.533196

**Authors:** Yilun Dong, Mingzhuo Li, Bryanna Cruz, Emily Ye, Yue Zhu, Lihua Li, Zhengjun Xu, De-Yu Xie

## Abstract

*Artemisia annua* is an effective antimalarial medicinal crop. We have established anthocyanin-producing red cell cultures from this plant with the overexpression of *Production of Anthocyanin Pigment 1* (*PAP1*) encoding a R2R3MYB transcription factor. To understand the molecular mechanism by which PAP1 activated the entire anthocyanin pathway, we mined the genomic sequences of *A. annua* and obtained eight promoters of the anthocyanin pathway genes. Sequence analysis identified four types of AC cis-elements from six promoters, the MYB response elements (MRE) bound by PAP1. In addition, six promoters were determined to have at least one G-Box cis-element. Eight promoters were cloned for activity analysis. Duel luciferase assays showed that PAP1 significantly enhanced the promoting activity of seven promoters, indicating that PAP1 turned on the biosynthesis of anthocyanins via the activation of these pathway gene expression. To understand how 2,4-dichlorophenoxyacetic acid (2,4-D), an auxin, regulates the PAP1-activated anthocyanin biosynthesis, five different concentrations (0, 0.05, 0.5, 2.5, and 5 μM) were tested to characterize anthocyanin production and profiles. The resulting data showed that the concentrations tested decreased the fresh weight of callus growth, anthocyanin levels, and the production of anthocyanins per petri dish. HPLC-qTOF-MS/MS based profiling showed that these concentrations did not alter anthocyanin profiles. Real time RT-PCR was completed to characterize the expression *PAP1* and four representative pathway genes. The results showed that the five concentrations reduced the expression levels of the constitutive *PAP1* transgene and three pathway genes significantly and eliminated chalcone synthase gene in expression either significantly or slightly. These data indicate that the constitutive *PAP1* expression depends on gradients added in the medium. Based on these findings, the regulation of 2,4-D is discussed for anthocyanin engineering in red cells of *A. annua*.

**Conclusion:** Promoters of eight anthocyanin pathway genes were cloned. Four types of AC cis-elements were identified from six promoters and G-Box elements were also determined from six promoters. PAP1 enhanced the activity of eight promoters. 2,4-D downregulated the expression of the constitutive PAP1 transgene leading to the decrease the biosynthesis of anthocyanins.

## Introduction

*Artemisia annua* is the natural source for the production of artemisinin, an effective antimalarial sesquiterpene lactone endoperoxide (Qinghao-Kanglue-Hezuo-Yanjiu-Zu 1977; Tu 2011). To date, artemisinin-based combination therapy (ACT) forms the frontline therapy of malarial diseases (Ashley and White 2005). Especially, ACT effectively treats severe malarial diseases caused by *Plasmodium falciparum,* which is resistant to other antimalarial medicines and leads to approximately 80% of malaria-associated deaths (Nosten and White 2007; WHO 2010, 2006). One main reason is the insufficient yield of artemisinin produced from the *A. annua cropping.* This long problem has resulted from the low and variable content of artemisinin in plants (Xie et al. 2016; Brown 2010). To solve this problem, numerous global efforts have been made to breed or engineer superior *A. annua* plants (Xie et al. 2016; Alejos-Gonzalez et al. 2011). Furthermore, a large number of phytochemical investigations have developed a fundamental platform to understand the complexity of metabolisms that can affect artemisinin biosynthesis. To date, isolation efforts have elucidated more than 600 compounds in *A. annua*, including monoterpenoids, sesquiterpenoids, triterpenoids, flavonoids, and other metabolites (Brown 2010). These metabolites characterize metabolic networks that can enhance metabolic engineering of artemisinin for high production.

Although no anthocyanin structures were reported from *A. annua* in the past (Brown 2010), we recently used Production of Anthocyanin Pigment 1 (PAP1), a master MYB regulator of Arabidopsis anthocyanin biosynthesis, to have established red cell cultures from this antimalarial plant (Judd et al. 2023). We detected 17 anthocyanin molecules from the extracts of red cells. More importantly, we identified two anthocyanidins and two anthocyanins, which were structurally characterized to be pelargonidin, cyanidin, pelargonidin-3-O-glucose, and cyanidin-3-O-glucose. Furthermore, the characterization of anthocyanidins indicate that cyanin and pelargonin are the main groups of anthocyanins in red cells perhaps in this antimalarial plant. In addition, the engineering of anthocyanins did not exclude the biosynthesis of artemisinin in red cells, indicating that although the formation artemisinin is solely proposed to occur in an oxidative condition, red cells engineered with an antioxidative subcellular condition possessed a new mechanism for the biosynthesis of artemisinin. Given that the production of anthocyanins in *A. annua* is anticipated to be useful to obtain value-added antimalarial plants, these findings are important to engineer *A. annua* for antimalarial application in the future. The reason is that anthocyanins are important nutraceuticals benefiting human health, such as antioxidative (Kano et al. 2005; Pool-Zobel et al. 1999; Denev et al. 2010; Shimizu et al. 2010; Tsuda et al. 1994), anti-cancer (Bowen-Forbes et al. 2010; Barrios et al. 2010; Dai et al. 2009; Faria et al. 2010), and neuron-protective properties.

To promote the use of red cells for high production and structural diversity of anthocyanins, it is critical to optimize the composition of tissue culture medium. In our previous metabolic engineering of anthocyanins in *Arabidopsis* and tobacco cells, we observed that nitrogen, carbon, and plant hormones could affect both cell growth and anthocyanin biosynthesis (Shi and Xie 2011; Zhou et al. 2012; Liu et al. 2014). In engineered red Arabidopsis cells developed from *pap1-D* mutant, the biosynthesis of anthocyanins was activated by the upregulation of *PAP1* driven by its natural promoter, to the front of which four 35S enhancers were inserted by T-DNA activation tagging (Shi and Xie 2011; Borevitz et al. 2000). We found that plant auxins, such as 2, 4-dichlorophenoxyacetic acid (2, 4-D), indole-acetic acid (IAA), and 1-naphthaleneacetic acid (NAA) could regulate the biosynthesis of anthocyanins in red *pap1-D* cells (Liu et al. 2014). In our engineered red *A. annua* cells, we used a 35S promoter to overexpress *PAP1* constitutively. Although whether auxins can regulate the biosynthesis of anthocyanin activated by the constitutive expression of *PAP1* driven by the 35S promoter is unknown, based on our previous data, we hypothesize that different concentrations might control the *PAP1* transgene expression, thus leading to alteration of the anthocyanin production.

In this investigation, we mined the promoters of eight anthocyanin pathway genes from the publically available genomic sequences and identified cis-elements that are bound by PAP1. In addition, G-Box cis-elements were identified from these promoter sequences. Dual luciferase assays were completed to understand the regulation mechanisms of PAP1 in red cells. Based on these data, we tested effects of 2,4-D on the biosynthesis of anthocyanins in the engineered red cells of *A. annua.* This auxin downregulated the overexpression of the *PAP1* transgene and decreased the biosynthesis of anthocyanins in red cells.

## Materials and methods

### Reagents

Most tissue culture gradients, including kinetin, 2, 4-D, sucrose, phytoagar, macronutrients, micronutrients and organic nutrients used in the MS medium were purchased from Plant Media (Dublin, OH, USA). Hydrochloric acid (36.5–38%) was purchased from BDH (cat# BHH3028-2.5L, Westchester, PA, USA). Organic solvents used for analytic experiments included acetonitrile (LC-MS grade) from EMD (cat# AX0156-1, Gibbstown, NJ 08027, USA) and acetic acid (HPLC grade, cat# 9515-03), and methanol (LC–MS grade, cat# 9830-03) purchased from J. T. Baker (Phillipsburg, NJ 08865, USA). The cyanidin, pelargonidin, and delphinidin standards were purchased from Indofine (Hillsborough, NJ, USA). Other reagents are specified in the experiments described below.

### Medium preparation and callus culture

We recently reported our methods and steps for medium preparation and subculture of red cells (Judd et al. 2023). Herein, we followed our protocols reported. In brief, we developed four types red cells, TAPA1, 2, 3, and 4. Since TAPA1 produced the highest contents of anthocyanins, this cell line was used to test effects of 2,4-D on anthocyanin biosynthesis. In addition, vector control (VC) cells were used as controls. The subculture medium was composed of basal MS medium (Murashige and Skoog 1962) supplemented with 1 mg/L NAA, 0.1 mg/L kinetin, and 1 mg/L of ascorbic acid, and 3% sucrose. The subculture period was 20 days. The lighting condition and temperature were set up 16/8 (light/dark) hours of photoperiod and 25°C, respectively. The light intensity was 50 μmol/min, m^2^.

### Tests of 2, 4-D treatments

To test effects of 2, 4-D on anthocyanin biosynthesis in red cells, the 1 mg/L NAA used in the subculture medium was replaced with five concentrations (0, 0.05, 0.5, 2.5, and 5.0 μM). Subculture medium was used as a control. Twenty milliliters of warm and melted agar medium were poured into each petri dish (15 × 100 mm, height × diameter) and solidified at room temperature. Six petri dishes were prepared for each concentration treatment. Each petri dish was inoculated with 0.3 g fresh weight of calli (TAPA1or VC cell lines) as one technical replicate. All petri dishes were placed in a tissue culture chamber. The growth condition was the same as described above. Calli were grown for 20 days and then harvested to measure fresh weights. The harvested calli were frozen in liquid nitrogen and stored in a −80°C freezer for other experiments described below. This experiment was repeated three times.

### Real time reverse transcription-polymerase chain reaction

Total RNA was isolated from the six types of calli, including VC, subculture red cells, 0.05 μM, 0.5 μM, 2.5 μM, 5 μM 2,4-D treated calli. One hundred mg of calli was ground into a fine powder in liquid nitrogen, which was placed in a 1.5 mL Eppendorf tube for total RNA extraction. The IBI isolate (IBI Scientific, USA) kit was used to extract total RNA. The extraction steps followed the manufacturer’s instructions. The extracted RNA samples were treated with DNase as described previously (Judd et al. 2023). The resulting DNA-free RNA samples were stored in a −80 freezer. The first strand cDNA was synthesized using the DNA-free RNA samples as template and the High-Capacity cDNA Reverse Transcription Kit (Applied Biosystems, USA). The reverse transcription steps followed the manufacturer’s instructions.

The iTaq-Universal SYBR Green Supermix was used for quantitative reverse transcription-polymerase chain reaction (qRT-PCR) by following the manufacturer’s instructions (Bio-Rad, USA). The first strand cDNA samples were diluted 3 times prior to qRT-PCR. The cDNAs of the *AtPAP1 transgene* and four representative anthocyanin pathway genes, *AaPAL1, AaCHS, AaDFR,* and *AaANS,* were amplified by qRT-PCR. The gene-specific primers are listed in supplementary table 1. The thermal cycle consisted of 50°C for 10 min, 95°C for 10 min, and 45 cycles of 95°C for 15 secs and 60°C for 1 min. Beta-ACTIN was used as an internal control for normalization. Three technical replicates were performed for each biological sample. In order to quantify the relative expression levels of each gene among different biological samples, the ΔΔCt algorithm values were calculated. For comparison, the relative expression levels of each gene were normalized to the vector control samples.

### Anthocyanin extraction and measurement

A total of 100 mg of frozen calli powder for each sample was suspended in 1 mL of extraction buffer (0.5% HCl in 100% methanol) contained in a 1.5 mL Eppendorf extraction tube. The methods for the extraction and quantification of anthocyanins were as described previously (Shi and Xie 2011). A minor modification was completed. In brief, 200 μL extract for each sample was pipetted to one clear well in a 96-well plate and then read absorption values at the wavelength of 530 nm on a high-performance microplate reader (SpectraMax® iD3). Three technical replicates were performed for each biological sample. The anthocyanin level in calli was indicated with absorption value. In addition, an authentic standard curve was developed with cyanidin 3-O-glucoside. The standard curve was used to calculate anthocyanin production per petri dish.

### HPLC–qTOF–MS analysis of anthocyanins

The profiles of anthocyanins were analyzed on Agilent 6210 time-of-flight LC/MS (Agilent Technologies, Santa Clara, CA, USA) as described previously (Judd et al. 2023). In brief, the HPLC mobile phase solvents consisted of 1% acetic acid in LC-MS grade water (A) and 100% acetonitrile (B). An Eclipse XBD-C18 analytical silica column (250 mm x 5 mm) was used for separation. A gradient system consisting of different A to B ratios was developed to elute anthocyanins and anthocyanidins. The ratios of solvent A to B and their elution time were composed of 90:10 (0–5 min), 90:10–88:12 (5–10 min), 88:12–80:20 (10–20 min), 80:20–75:25 (20–30 min), 75:25–65:35 (30–35 min), 65:35–60:40 (35–40 min), 60:40–50:50 (40–55 min), and 50:50–10:90 (55–60 min). The column was washed for 10 min with 10% solvent B after each sample. The injection volume was 10 μL. The flow rate was 0.4 mL/ min. A photodiode array detector was used to record metabolites at 530 nm. The drying gas flow was set to 12 L/min, and the nebulizer pressure was set to 35 psi.

### Cloning of promoters from *Artemisia*

Promoter sequences of eight genes *(AaPAL, AaC4H, Aa4CL2, AaCHS, AaCHI2, AaF3H, AaDFR,* and *AaANS*) involved in the flavonoid pathway were identified from genomic sequences of *A. annua* (http://www.ncbi.nlm.nih.gov/) in order to understand the regulatory function of *PAP1* in transgenic cells. Each sequence was comprised of 1.6 kb of nucleotides located in the immediate upstream of the coding region. AC elements were identified using the PLACE approach (https://www.dna.affrc.go.jp/PLACE/?action=newplace). Based on the sequences, promoters were cloned from *A. annua* as described before (Li et al. 2022). Forward (5’ to 3’ end) and reverse (3’-5’) primer pairs (Supplementary Table S2) were designed to amplify at least 1 kb of nucleotides located immediately the upstream of the coding regions of the eight genes. The amplified genomic DNA fragments were purified with a gel purification kit (Qiagen) as described previously (Li et al. 2022). All the eight purified promoter fragments were cloned into the pEASY-T1 plasmid and confirmed by sequencing as described previously (Li et al. 2022). After nucleotides of all the sequences were confirmed correctly, they were used for dual luciferase assay described below.

### Development of reporter and effector constructs

Eight reporter constructs were developed for dual luciferase assay described below. The promoters of *AaPAL, AaC4H, Aa4CL2, AaCHS, AaCHI2, AaF3H, AaDFR* and *AaANS* were used to develop reporter vectors. We cloned these promoters into the pGreenII-0800 vector to develop reporter constructs as described previously (Li et al. 2022). In brief, the pGreenII-0800 vector contains multiple restriction enzyme sites immediately at the N-terminus of a firefly-luciferase gene. To clone promoters into the pGreenII-0800 vector, primers were designed to add PstI and Smal I restriction sites in the two ends of each promoter sequence (Supplementary Table S2). PCR was completed to amplify each promoter sequence with the same thermal programs (Supplementary Table S2), which was purified as described above. pGreenII 0800 and promoters of *AaPAL, AaC4H, Aa4CL2, AaCHS, AaCHI2, AaF3H, AaDFR* and *AaANS* were digested with PstI and Smal I and then ligated using T4 ligase. The resulting ligation products were transformed into competent cells Top10. Positive *E. coli* colonies were obtained from colony screening on LB medium supplemented with 50 mg/ml kanamycin. The positive colony strains were screened by PCR and then confirmed by sequencing. As a result, eight reporter constructs, namely as *AaPAL-, AaC4H-, Aa4CL2-, AaCHS-, AaCHI2-, AaF3H-, AaDFR-* and *AaANS-* pGreenII-0800 (Fig. 3 a), were obtained for dual luciferase assays described below.

**Fig. 1.**
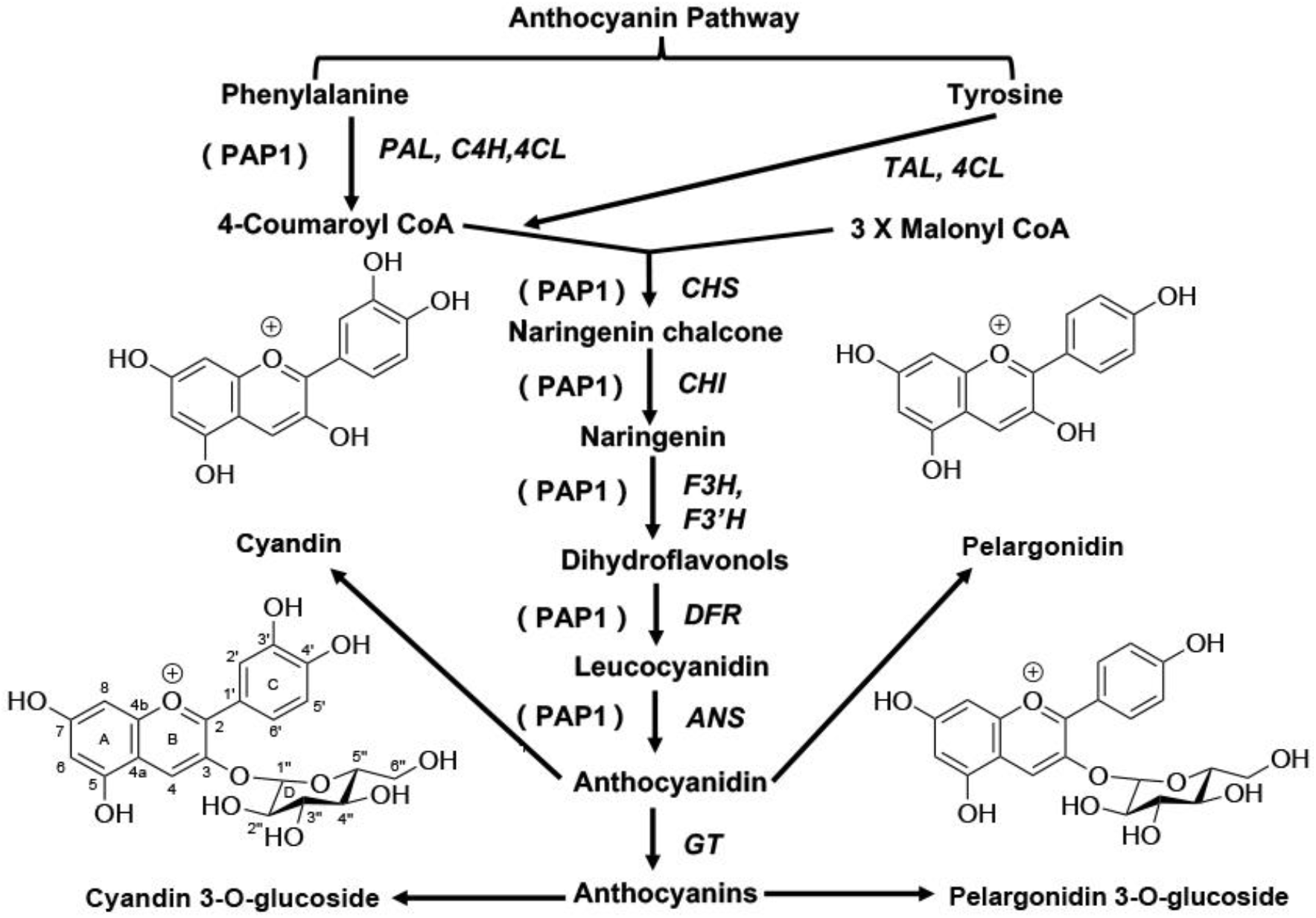
The biosynthetic pathway of anthocyanins activated by PAP1 in engineered red cells of *A. annua.* The biosynthetic pathway of anthocyanins were recently characterized in engineered TAPA cells of *A. annua* (Judd et al 2023). Cyanins and pelargonins are two groups of anthocyanins produced in TAPA cells. The pathway genes include the beginning steps genes, *PAL*: phenylalanine ammonia lyase, *TAL*: tyrosine ammonia lyase, *C4H:* cinnamate-4-hydroxylase and *4CL*: 4-coumaryol CoA ligase; the early genes, *CHS*: chalcone synthase, *CHI*: chalcone isomerase, *F3H:* flavanone-3 hydrolase and *F3’H:* flavonoid-3’-hydroxylase; and the late genes, *DFR*: dihydroflavonol reductase, *ANS:* anthocyanidin synthase (also called *LDOX*: leucoanthocyanidin dioxygenase) and *GT:* glycosyltransferase. *PAP1:* production of anthocyanin pigmentation 1.

**Fig. 2.**
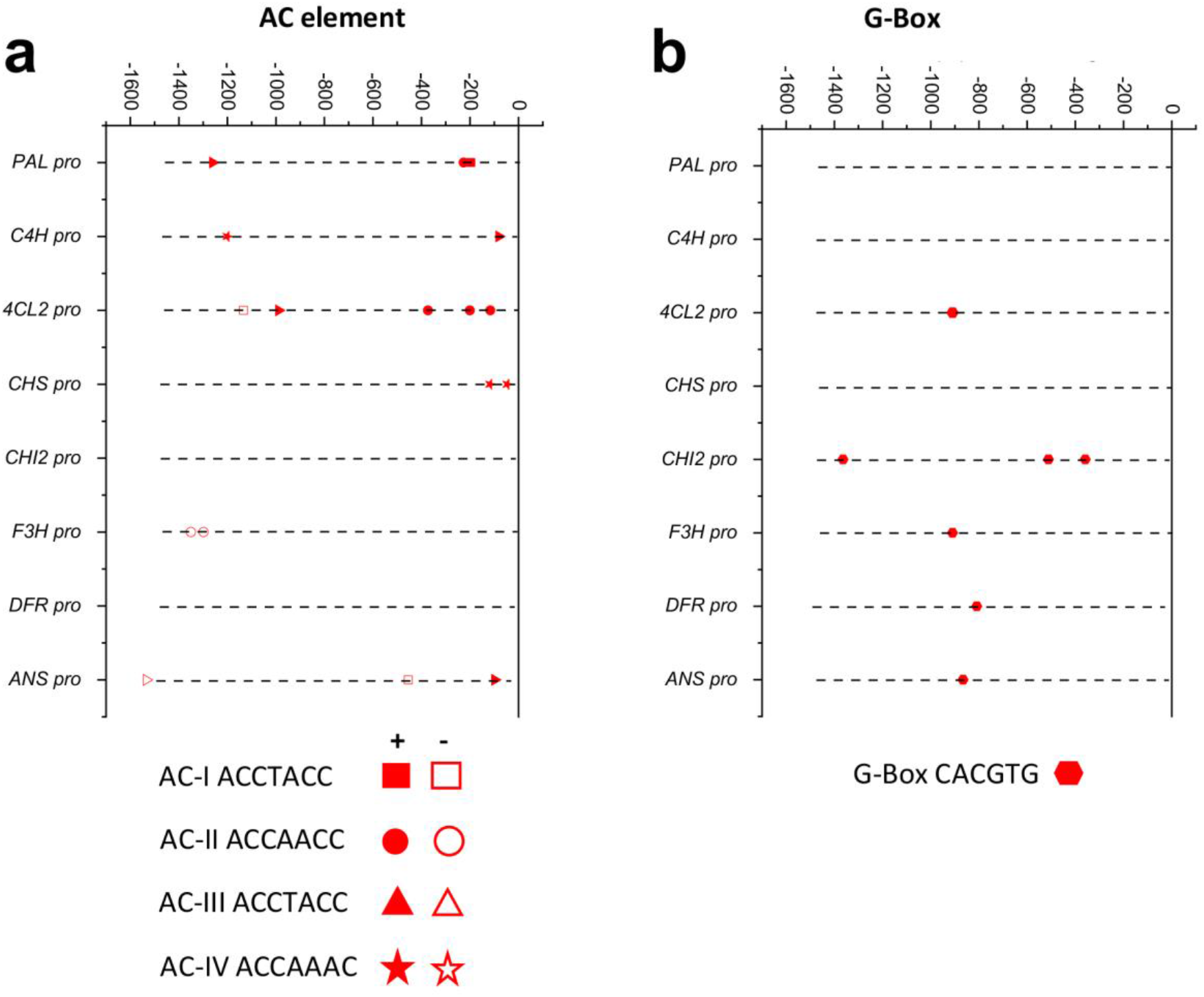
AC and G-Box cis-elements identified in the promoters of anthocyanin pathway genes. a, four types of AC cis-elements are identified in the promoters of six pathway genes. the direction from 5’ to 3’ and “-”: the direction from 3’ to 5’. b, G-Box cis-elements are identified in the promoters of six pathway genes.

**Fig. 3.**
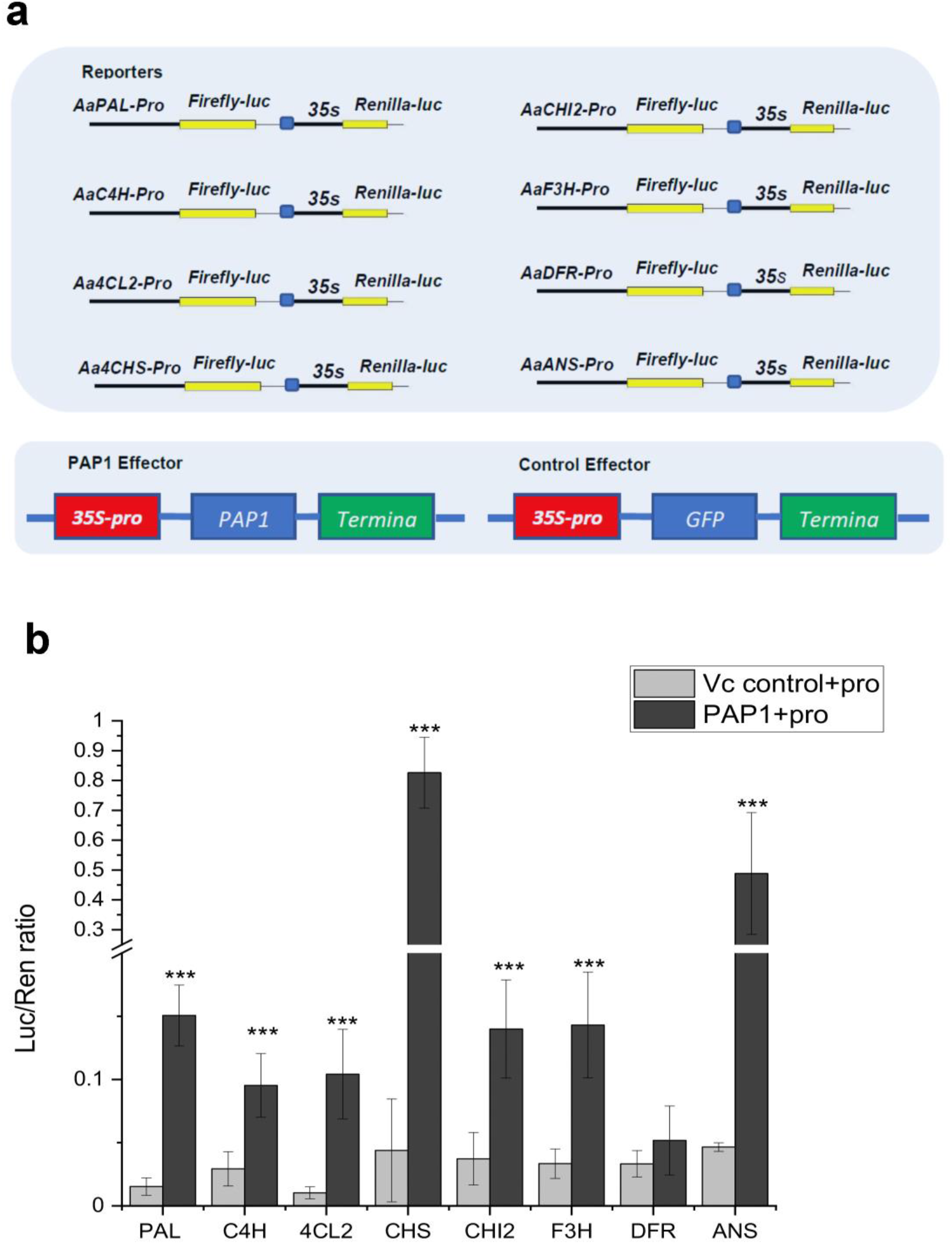
Results of dual luciferase assays. a, cassettes show eight reporter vectors that are developed with eight promoters fused with a firefly luciferase gene, one cassette exhibits an effector vector developed from PAP1, and one cassette shows GFP as a control effector. b, Luciferase activity data show that PAP1 significantly enhances the activity of seven promoters and slightly increases the activity of the DFR promoter. Bars labeled with *** means the number significantly higher from PAP1 than from GFP effector controls (p-value < 0.01).

Meanwhile, the ORFs of *GFP* and *PAP1* were cloned into the effector vector PK2GW7, in which each was driven by a 35S-promoter as described previously (Li et al 2022). In brief, gateway cloning was performed to develop two effector vectors. PCR was performed to amplify each ORF with an attB in the 5-end and an attB2 adapter in the 3-end. Primers are listed in supplementary table S2. Each PCR product was purified and then ligated into the pDonr207 plasmid with BP reactions, which followed the protocol of Gateway BP Cloning Reaction Kit. Then, the *GFP* and *PAP1* in the pDonr207 plasmid were cloned to the expression vector PK2GW7 via LR reaction by following the instruction of the Gateway LR Cloning Reaction Kit. The resulting recombinant PK2GW7*-*GFP and PK2GW7-PAP1 plasmids (Fig. 3 a) were introduced to *E. coli* Top10 to select positive colonies on an LB medium containing 50 mg/l spectinomycin as described above.

### Dual luciferase assay

*Agrobacterium tumefaciens* GV3101was used for transient expression of both effectors and reporters. Each of *AaPAS-pro AaC4H-pro, Aa4CL2-pro, AaCHS-pro, AaCHI2-pro, AaF3H-pro, AaDFR-pro,* and *AaANS-pro*-pGreenII-0800 vectors and a cooperative enhancer vector pSoup+p19 were co-transformed into competent GV3101 cells through electroporation shock method with 2000V. Meanwhile, two effector vectors, *GFP*-PK2GW7 and *PAP1*-PK2GW7, were transformed into competent *Agrobacterium* GV3101, respectively. Selection steps for positive antibiotic-resistant recombinant GV3101 colonies followed our protocol reported previously (Li et al. 2022). As a result, eight reporter and two effector GV3101 colonies were obtained for dual luciferase assay.

Each of eight reporter and two effecter colonies was inoculated to 50 mL fresh liquid LB medium supplemented with antibiotics contained in a 250 ml E-flask. Each flask was placed on the same shaker with a speed of 120 rpm to culture for 12 hours, when the optical density values (OD600) of suspension cultures were 0.6 recorded at the wavelength of 600 nm. Cultured *Agrobacterium* cells were harvested to a 50 mL tube and centrifuged at 4000 rpm for 10 min. The resulting *Agrobacterium* pellet was suspended in infiltration buffer (10mM pH5.5 MES, 10mM MgCl2, 150 μM acetosyringone) and the concentrations were adjusted to an OD600 value of 0.2-0.3. The tubes containing the suspended *Agrobacterium* cells were placed on the same shaker and incubated for 2 hrs at room temperature prior to transient expression experiments.

Transient expression assay was performed in leaves of *Nicotiana benthamiana* using activated *Agrobacterium* GV3101 cells. 100 μl of reporter *Agrobacterium* GV3101 cells (each of eight promoters) and 450 μl effecter *Agrobacterium* GV3101 cells (PAP1 or GFP) were mixed appropriately. As a result, eight paired mixtures were formed for infiltration. Meanwhile, the empty pGrenn-II-0800 (reporter) and *GFP* gene effecter were formed a pair as a control. Then, 200 μl of the each mixed culture was infiltrated into three locations on each of four young leaves from 35-day old *N. benthamiana,* each leaf as one technical replicate. Five independent plants were infiltrated for each paired effector and reporter as five biological replicates. After infiltration, all plants were grown in the phytotron for four days prior to analysis of dual-luciferase as described below.

Dual-luciferase assay was completed using a Dual-Luciferase Reporter Assay System kit (Promega, Madison, USA) by following the manufacturer’s instructions as described previously (Li et al. 2022). In brief, leaf discs with a 1.0 cm diameter of infiltration were punched into a 1.5 ml tube and then homogenized in 500 μl Passive Lysis Buffer. The homogenized mixture was centrifuged for 10 min at 12,000 rpm at 40°C. The resulting supernatant was transferred to a new 1.5 mL tube and then diluted 50 times with the Passive Lysis Buffer. Ten μl diluted extract was mixed with 40 μl Luciferase Assay Buffer. The firefly luminescence (LUC) from samples was immediately recorded for 5.0 sec on a GloMax 20/20 luminometer (Promega, Madison, USA) followed by 10-sec interval. Then, the reaction was stopped by adding 40 μl Stop and Glow Buffer and immediately recorded for a second luminescence using renilla (REN) luminescence. The ratios of LUC to REN were calculated. For each pair of a reporter (promoter) and an effector (PAP1), five biological replicates were performed to obtain LUC to REN ratio values. In addition, five biological replicates were performed for eight reporters and *GFP* effecter as controls.

### Statistical analysis

Student’s t test was performed to analyze the significant differences in dual luciferase assays. One way ANOVA was used to statistically compare calli fresh weights, total anthocyanins and gene expression.

## Results

### Characterization of AC and G-box cis-elements in promoters of anthocyanin pathway genes of *A. annua*

PAP1 activates the biosynthesis of anthocyanins via binding to AC-elements of promoters (Li et al. 2022). To understand the regulation mechanisms of anthocyanin biosynthesis in PAP1-based *A. annua* cells, we mined the genomic sequences of this antimalarial plant (Shen et al. 2018; Liao et al. 2022). The nucleotide sequences of promoters were identified in the upstream of open reading frame (ORF) of *AaPAL*, *AaC4H*, *Aa4CL2*, *AaCHS*, *AaCHI2*, *AaF3H*, *AaDFR*, and *AaANS* (Supplementary materials). The length for all of them was 1.6 kb. Sequence analysis identified that the promoters of *AaPAL*, *AaC4H*, *Aa4CL2*, *AaCHS*, *AaF3H*, and *AaANS* contained 2, 2, 5, 3, and 3 AC cis-elements, respectively, while no AC-elements were found in the promoters of *AaCHI2* and *AaDFR* (Fig. 2 a and Supplementary materials). The total 15 AC elements include four types, AC-I: ACCTACC, AC-II; ACCAACC; AC-III: ACCTACC, and AC-IV: ACCAAAC. Further analysis revealed that the four types had two directions located in the genome, either 5’ to 3’ or 3’to 5’ (Fig 2 a and Supplementary materials). The promoters of *AaPAL, AaC4H*, and *AaCHS* have AC-I and II, AC-III and IV, and two AC-IV elements, respectively, which are featured by the 5’ to 3’ direction. The promoter of *Aa4CL2* has one AC-I element with the 3’to 5’ direction, three AC-II elements with the 5’ to 3’ direction, and one AC-III element with the 5’ to 3’ direction. The promoter of *AaF3H* has two AC-II elements featured by the 3’ to 5’ direction. The promoter of *AaANS* has one AC-I from 5’ to 3’ and two AC-II elements one from 5’ to 3’ and the other from 3’ to 5’.

PAP1 partners TT8 and TTG8 homologs form a MBW complex to activate the biosynthesis of anthocyanins and TT8 binds to a G-box element in plants (Li et al. 2022; Shi and Xie 2011). To understand G-box features, we analyzed the eight promoter sequences. The resulting data showed that the promoters of *Aa4CL2, AaF3H, AaDFR,* and *AaANS* had one G-Box element and that of AaCHI2 has one, while *AaPAL, AaC4H,* and *AaCHS* did not have (Fig. 2 b and Supplementary Materials).

#### PAP1 increases the promoting activity of seven promoters

Duel luciferase assay was completed to understand the regulatory mechanism of PAP1 in engineered TAPA1 cells. One kb long promoter sequences of *AaPAL*, *AaC4H*, *Aa4CL2*, *AaCHS*, *AaCHI2*, *AaF3H*, *AaDFR*, and *AaANS* were amplified from our self-pollinated *A. annua* plants. Further sequence analysis confirmed the presence of AC cis-elements or G-Box elements in the promoters as described above. Eight promoters were fused with a luciferase gene in the pGreenII 0800 vector to obtain eight reporter constructs and *PAP1* was cloned to the PK2GW7 vector to obtain a PK2GW7-PAP1 effector (Fig. 3 a). In addition, PK2GW7-GFP was constructed as a control effector. The PK2GW7-PAP1 effector and the eight reporter vectors formed eight pairs for transient expression in leaves of *N. benthamiana.* In addition, the PK2GW7-GFP vector and the eight report vectors formed eight control pairs. After transient expression, the luciferase analysis showed that the activities of the *AaPAL, AaC4H, Aa4CL2, AaCHS, AaCHI2, AaF3H,* and *AaANS* were significantly increased compared with controls (Fig. 3 b). The promoting activity of the *AaDFR* promoter was slightly increased. These data show that PAP1 activates the biosynthesis of anthocyanins in TAPA1 cells via the activation of the promoters of main pathway genes.

#### Effects of 2,4-D on anthocyanin production

Five concentrations of 2, 4-D were tested to understand its effect on PAP1-activated anthocyanin formation in TAPA1 cells. After inoculation of cells, the growth of calli was photographed every 3 or 5-day to observe pigmentation phenotypes and biomass increases (Fig. 4). Direct comparison of calli growth indicated that the biomass of TAPA1 calli exhibited a reduction trend as the concentrations of 2, 4-D were increased from 0 to 5 μM. Pigmentation comparison indicated that the red pigmentation of anthocyanins was not obviously altered on different media (Fig. 4). Vector control calli did not produce pigments. The measurement of fresh weight indicated a reduction of trend of calli biomass per petri dish as the concentrations were increased from 0 to 5 μM (Fig. 5 a). The biomass of vector control calli exhibited a reduction trend. These data indicate that high 2,4-D concentrations inhibited callus growth. The absorption measurement of total anthocyanins indicated that the levels of anthocyanins were diminished by all five concentrations. Cyanidin-3-O-glycoside was used as a standard to estimate the production of anthocyanin from each petri dish as this standard equivalent. The results showed that the production per petri dish was reduced. It was interesting that in the five concentrations tested, calli produced the most level of anthocyanins (Fig. 5 a). These data indicate that 2,4-D down-regulates the biosynthesis of anthocyanins engineered in TAPA1 cells.

**Fig. 4.**
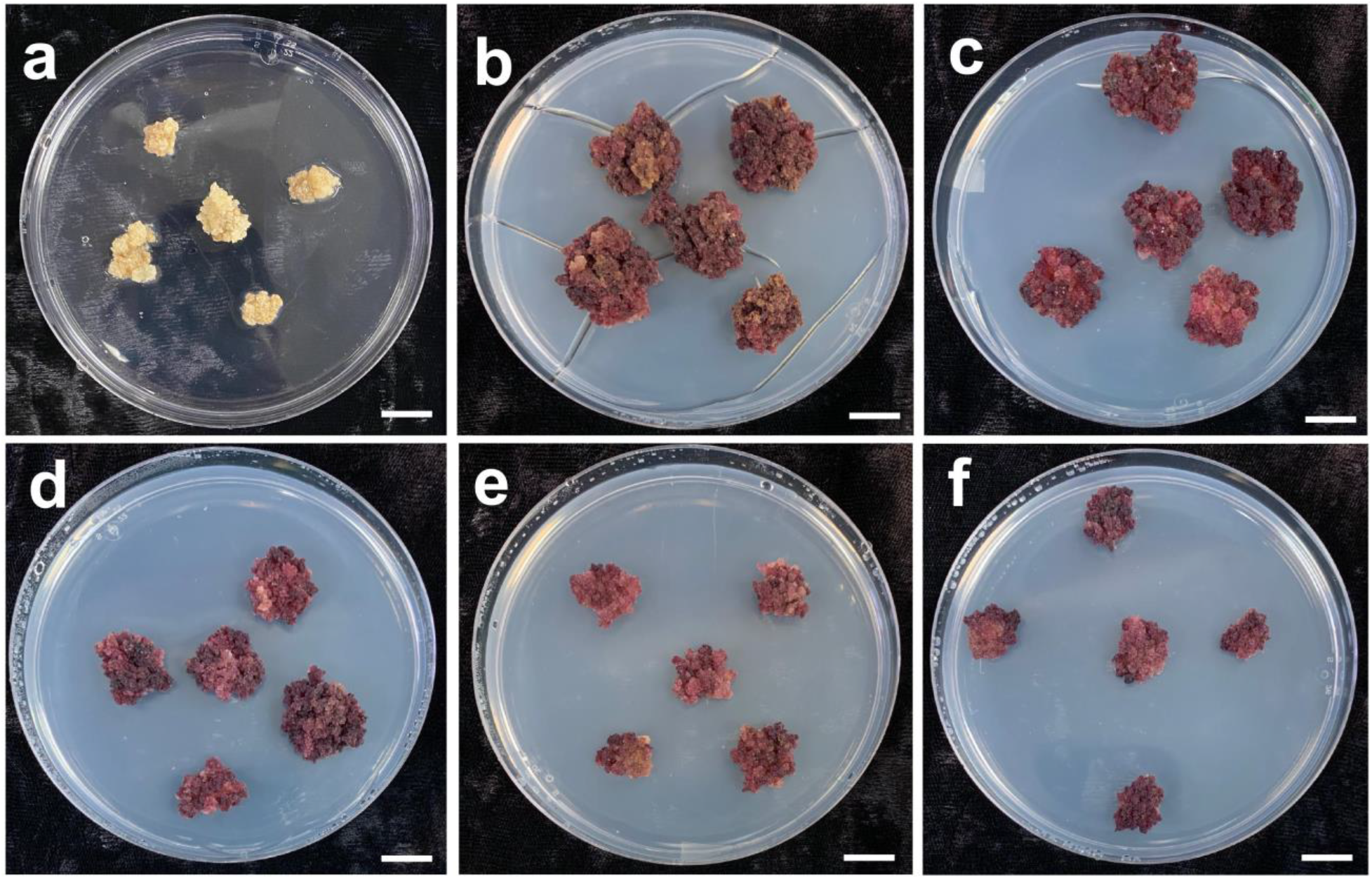
Phenotypes of red TAPA1 calli grown on five media supplemented with five 2,4-D concentrations. a, vector control calli grown on the medium without 2,4-D and b-f, TAPA1 calli grown on media containing 0 (b), 0.05 (c), 0.5 (d), 2.5 (e), and 5.0 (f) μM. The initial inoculum was 0.3 g fresh weight calli per petri dish. The photos were taken after 20-day of culture.

**Fig. 5.**
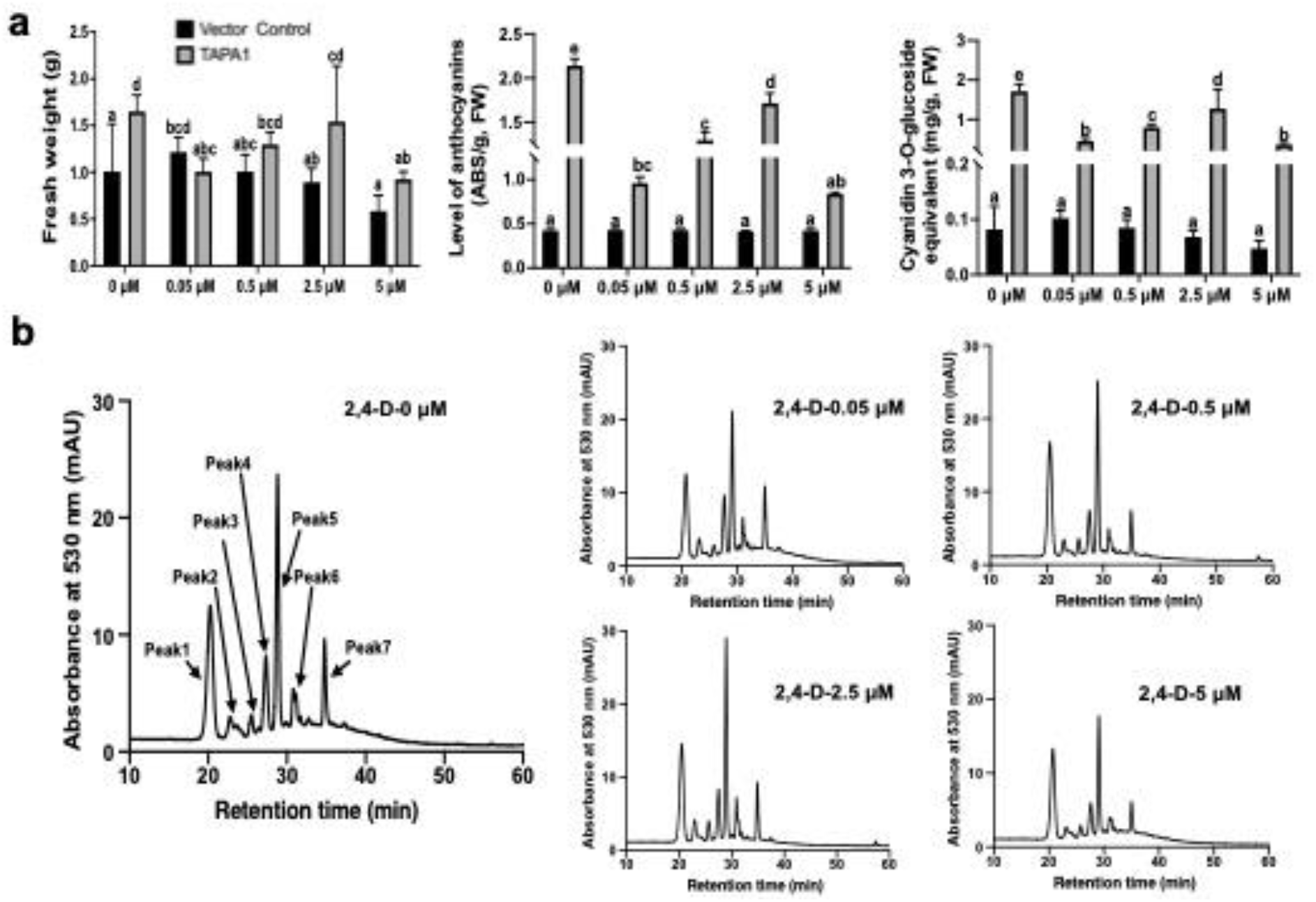
Effects of five different concentrations of 2, 4-D on the growth of calli and anthocyanin formation. a, effects of 2, 4-D on fresh weights of calli, anthocyanin levels, and production per petri dish. The bars labeled with different letters represent significant difference (P value < 0.05), while those with the same letter are not significantly different (P value > 0.05). The production per petri dish was estimated as cyanidin 3-O-glycoside equivalent. b, HPLC chromatograms were recorded at 530 nm to compare anthocyanin peak profiles from TAPA1 calli grown on media supplemented with five different 2,4-D concentrations.

#### Effects of 2,4-D on anthocyanin profiles

We previously reported 17 anthocyanin peaks detected in TAPA cells (Judd et al. 2023). To understand whether 2, 4-D could alter the profiles of anthocyanins, HPLC-qTOF-MS/MS was performed. All peak profiles were recorded at 530 nm. The resulting data showed that the peak profiles were not altered by different 2, 4-D concentrations (Fig. 5 b). In comparison, the sizes of all peaks were slightly increased by 2.5 μM 2, 4-D but slightly decreased by 5 μM 2, 4-D. These data supported the measurement results described above.

#### Effects of 2, 4-D on expression of anthocyanin pathway genes

We recently reported that PAP1 activated the expression of anthocyanin pathway genes in TAPA cells (Judd et al. 2023). To understand the effects of 2, 4-D concentrations on the expression of pathway genes, qRT-PCR was completed to analyze *PAP1*, *AaPAL*, *AaCHS*, *AaDFR*, and *AaANS.* The resulting data showed that the expression level of PAP1 exhibited a decrease trend as the concentrations of 2, 4-D increased (Fig. 6). Corresponding to the reduction of *PAP1* expression, the expression levels of *AaPAL, AaDFR,* and *AaANS* exhibited a reduction trend too (Fig. 6). The expression level of *AaCHS* was also reduced by 2, 4-D. It was interesting that in five concentration tested, the expression level of *AaCHS* was higher at 2.5 μM than at other concentrations (Fig. 6 c). This datum supported the anthocyanin production was higher on the medium supplemented with 2.5 μM than with 4 other concentrations.

**Fig. 6.**
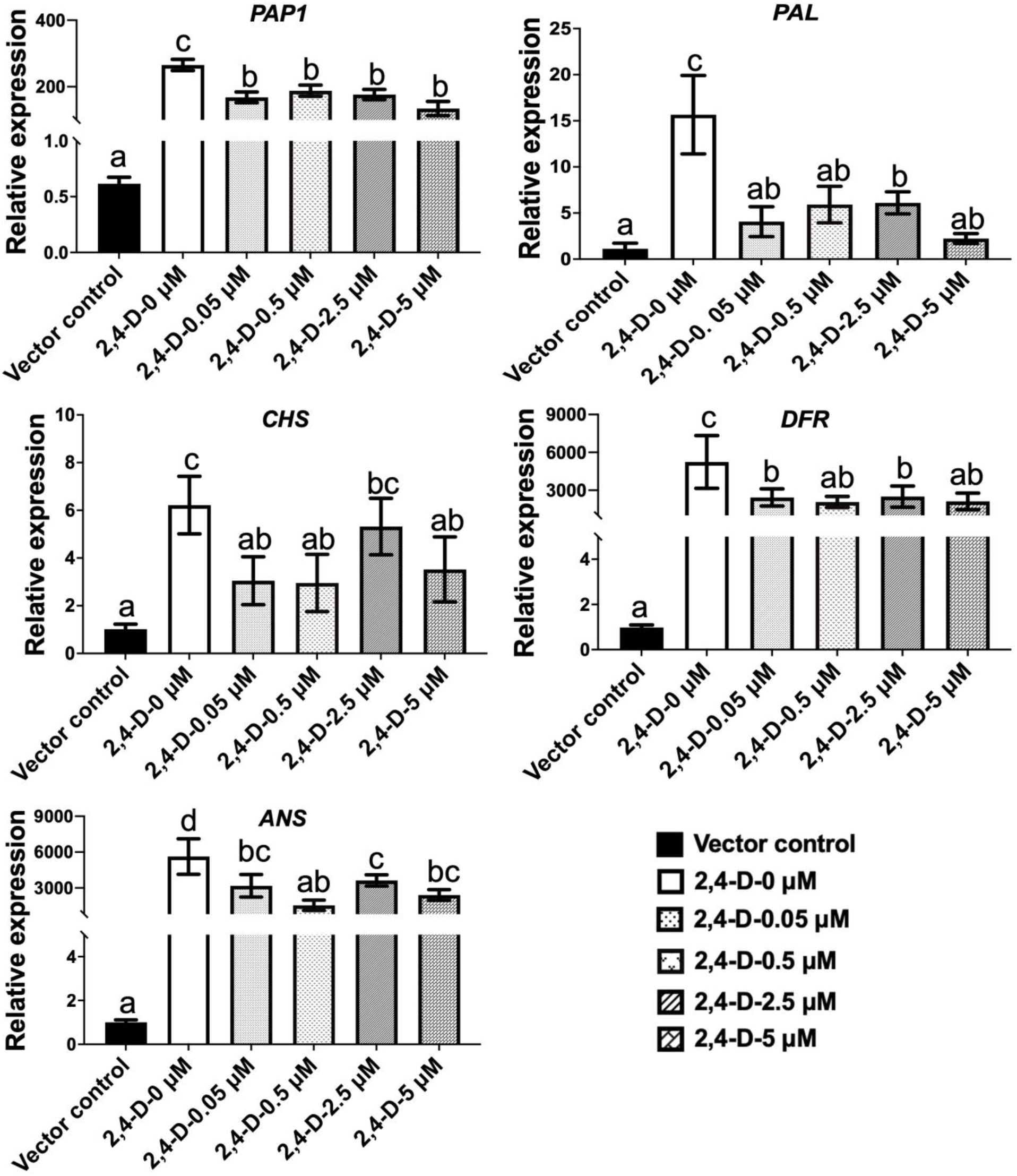
Effects of five different concentrations of 2, 4-D on gene expression in TAPA1 cells. The relatively expression levels of *PAP1, PAL, CHS, DFR,* and *ANS* were measured with qRT-PCR assays.

## Discussion

PAP1 has been used to engineer anthocyanins in plant cells and crops for new traits. As well characterized, *PAP1* encodes MYB75 that is a master regulator and partners with TT8/GL3 and TTG1 to form a MBW complex upregulating the anthocyanin biosynthesis in *A. thaliana* (Xie and Shi 2012; Zhou et al. 2012; Shi and Xie 2011). Although whether PAP1 needs its MBW complex to perform the regulatory activity in an overexpressed plant remains still open for answers, the past investigations have reported interesting results that the regulatory activity of the overexpressed *PAP1* depended upon different plants. On the one hand, one report showed that the overexpression of *PAP1* could not activate the biosynthesis of anthocyanins in alfalfa (Peel et al. 2009). On the other hand, other reports showed that the ectopic expression of *PAP1* activated the biosynthesis of anthocyanins in tobacco (Borevitz et al. 2000; Xie et al. 2006; He et al. 2017), hop (Gatica-Arias et al. 2012), and tomato (Butelli et al. 2008; Zuluaga et al. 2008). Based on our recent findings, the different consequences of the overexpression of *PAP1* likely resulted from whether the host plants could provide endogenous bHLH and WD4 homologs to form a functional MBW complex. We found that PAP1 bound to different types of AC cis-elements and its regulatory activity was enhanced by the formation of its MYB complex (Li et al. 2022). It was particularly interesting that its ectopic expression in tobacco plants activated the expression of its partner the endogenous *TT8* homolog *(NtTT8)* in transgenic plants (Li et al. 2022), likely leading to the formation of a hybrid PAP1/NtTT8/NtTTG1 complex. Accordingly, our observations not only interpreted why PAP1 could activate the biosynthesis of anthocyanin in all tissues of three tobacco genotypes (He et al. 2017; Li et al. 2022), but also suggested that alfalfa might not provide an endogenous TT8 homolog to partner with PAP1 to form a hybrid MYB complex for regulation. We recently reported to use *PAP1* to engineer four types of red TAPA cells that biosynthesized anthocyanins (Judd et al. 2023). Herein, our experimental data further provide evidence to understand the molecular mechanism by which PAP1 activates the biosynthesis of anthocyanins. The promoter sequence analysis identified AC cis-elements in the promoters of eight pathway genes. Dual luciferase assays demonstrated that PAP1 enhanced the promoting activity of the eight promoters. Although we have not analyzed endogenous *AaTT8* homologs, based on our previous report of tobacco *TT8* activation by PAP1 (Li et al. 2022), we hypothesize that PAP1 activates the *AaTT8* expression and then forms a hybrid PAP1-AaTT8-AaTTG1 homolog. The indirect evidence is that although the promoters of *CHI2* and *DFR* lack an AC-element (Fig. 2 a), their transcription is activated in TAPA1 cells (Fig. 6) (Judd et al. 2023). The upregulation of *CHI2* and *DFR* likely results from the activation of AaTT8 expressed in TAPA cells, given that they have G-Box elements (Fig. 2 b), to which AaTT8 bind. Since these data are the first report, it is anticipated that continuous investigations will enhance elucidating the regulatory mechanisms of PAP1 in *A. annua* plants.

TAPA1 cells provide a platform to understand whether or not 2,4-D can regulate anthocyanin biosynthesis activated by the constitutive expression of *PAP1* driven by a 35S promoter. As well known, 2, 4-D has been reported to negatively regulate the biosynthesis of anthocyanins and other plant secondary metabolites in non-transgenic cells (Ozeki and Komamine 1985; Robbins et al. 1996; Ozeki and Komamine 1986; Makunga et al. 1997; Zhao et al. 2014). To date, the regulation of 2,4-D on anthocyanin biosynthesis has gained appropriate interpretation at transcriptional level that genes are transcribed by their native promoters. Callus and cell suspension culture systems have been a good platform to understand the engineering of anthocyanin biosynthesis regulated by 2,4-D for a few decades (Ball 1967; Stickland and Sunderland 1972; Wellmann et al. 1976; Dougall et al. 1980; Al Qurraan et al. 2012; Asano and Otobe 2011). Example models of callus and cell suspension cultures include carrot (Dougall et al. 1980; Dougall et al. 1983; Rose et al. 1996; Hirner and Seitz 2000; Narayan et al. 2005; Ozeki et al. 2000), grape (Bao Do and Cormier 1991; Do and Cormier 1991a; Do and Cormier 1991b; Cormier et al. 1992; Yousef et al. 2004), and *Catharanthus roseus* (Hall and Yeoman 1986b, a). Based on those previous reports, the regulation of 2, 4-D can be summarized to two types. One is that the presence of 2,4-D tested in the culture medium highly inhibits the formation of plant natural products (Pala et al. 2022; Liao et al. 2018). Callus culture or cell-suspension culture experiments have shown that the removal of 2, 4-D from the media or a low concentration of 2, 4-D is necessary for the induction of anthocyanins (Ozeki and Komamine 1985; Yoshihiro and Atsushi 1985; Ozeki and Komamine 1986; Ozeki et al. 1990). 2, 4-D downregulates gene expression, leading to reductions in pathway enzymes, such as PAL and CHS, in carrot cell and callus cultures (Ozeki et al. 1990; Ozeki et al. 2000). The other is that an optimized concentration of 2,4-D can increase the production of plant natural products (Shi and Xie 2011; Liu et al. 2014). When the concentrations are higher than the optimized one, 2,4-D inhibits the biosynthesis of plant natural products (Meyer and Staden 1995). Previous investigations on anthocyanin biosynthesis in suspension-cultured carrot cells (Ozeki and Komamine 1986; Narayan et al. 2005; Ozeki and Komamine 1985), *Oxalis linearis* (Meyer and Vanstaden 1995), and strawberry (Mori et al. 1994) have reported that high levels of 2, 4-D strongly inhibit anthocyanin formation. Although the mechanisms behind the inhibition of plant secondary metabolism by 2,4-D remains elusive, apposite progress has been made at the molecular levels. We previously engineered anthocyanin-producing cells from Arabidopsis *pap1-D* mutant, which was generated by T-DNA activation tagging. This dominant mutant plant line contains four cauliflower mosaic virus 35S enhancers that are inserted to the region immediately adjacent to the promoter of *PAP1.* This insertion results in the high expression of *PAP1,* which leads to the high anthocyanin accumulation in most tissues. The past genetic and molecular investigations have characterized that PAP1 is a master regulator (Xie et al. 2006; Solfanelli et al. 2006; Zhou et al. 2008; Gonzalez et al. 2008; Zvi et al. 2012; Shi and Xie 2011; Zhou et al. 2012) and partners with bHLH (*TT8*, *transparent testa 8; TT2, transparent testa 2; GL3, glabra 3;* and *EGL3, enhancer glabra 3*) and WD40 *(TTG1, transparent testa glabra 1*) to form MBW complexes (Ramsay and Glover 2005; Zhang et al. 2003; Gonzalez et al. 2008; Shi and Xie 2010; Shi and Xie 2011; Xie and Shi 2012). We used this plant to engineer red *pap1-D* cells, which allowed us to characterize the regulatory complexes involved in anthocyanin biosynthesis regulated by 2, 4-D, NAA, and IAA (Liu et al. 2014). Of the three types of auxins, 2, 4-D showed the strongest inhibitive effects on the biosynthesis of anthocyanins. In the tested concentration range of 2.2-27 μM, 2, 4-D strongly inhibited the biosynthesis of anthocyanins. Mechanistic studies revealed that 2, 4-D inhibited the transcription of *PAP1,* thus perturbed the formation of the PAP1-T8/GL3-TTG1 (MBW) complex, which led to the transcriptional reduction of pathway genes, such as *CHS, DFR* and *ANS.* These results indicate that 2, 4-D inhibits the promoting activity of the *PAP1* promoter although four 35S enhancers are inserted. These results showed that 2,4-D negatively regulated activity of the promoter of *PAP1* in red *pap1-D* cells. However, these data do not provide evidence to understand whether 2,4-D regulates the expression of *PAP1* driven by a constitutive 35S promoter. Herein, our data showed that although *PAP1* was overexpressed by a 35S promoter, 2,4-D in the tested range 0.05-5 μM could downregulate its transcription (Fig. 6). Real time RT-PCR data indicated that four tested pathway genes, *PAL*, *CHS*, *DFR*, and *ANS*, were downregulated in expression in red TAPA1 cells (Fig. 6). It was interesting that the expression level of *CHS* was slightly decreased by 2.5 μM 2,4-D. Corresponding to these transcriptional reductions, the levels of total anthocyanins were also decreased by 2,4-D (Fig. 5 a). The production of anthocyanins per petri dish was also reduced due to the decrease of both callus biomass and anthocyanin contents (Fig. 5 a). These data indicate that 2,4-D in a range of concentration range can inhibit the biosynthesis of anthocyanins activated by the constitutively expressed PAP1. Based on these data, a model is proposed to enhance understanding the regulation of 2,4-D (Fig. 7). In TAPA1 cells, 2,4-D downregulates the transcription of the overexpressed *PAP1,* which leads to the reduction of the MYB75 level and transcriptional activity. Accordingly, the expression of main pathway genes are subsided, which lead to diminish the biosynthesis of anthocyanins. This model shows that although a 35S promoter is used to overexpress PAP1, the presence of 2,4-D can inhibit its regulation and diminishes the biosynthesis of anthocyanins.

**Fig. 7.**
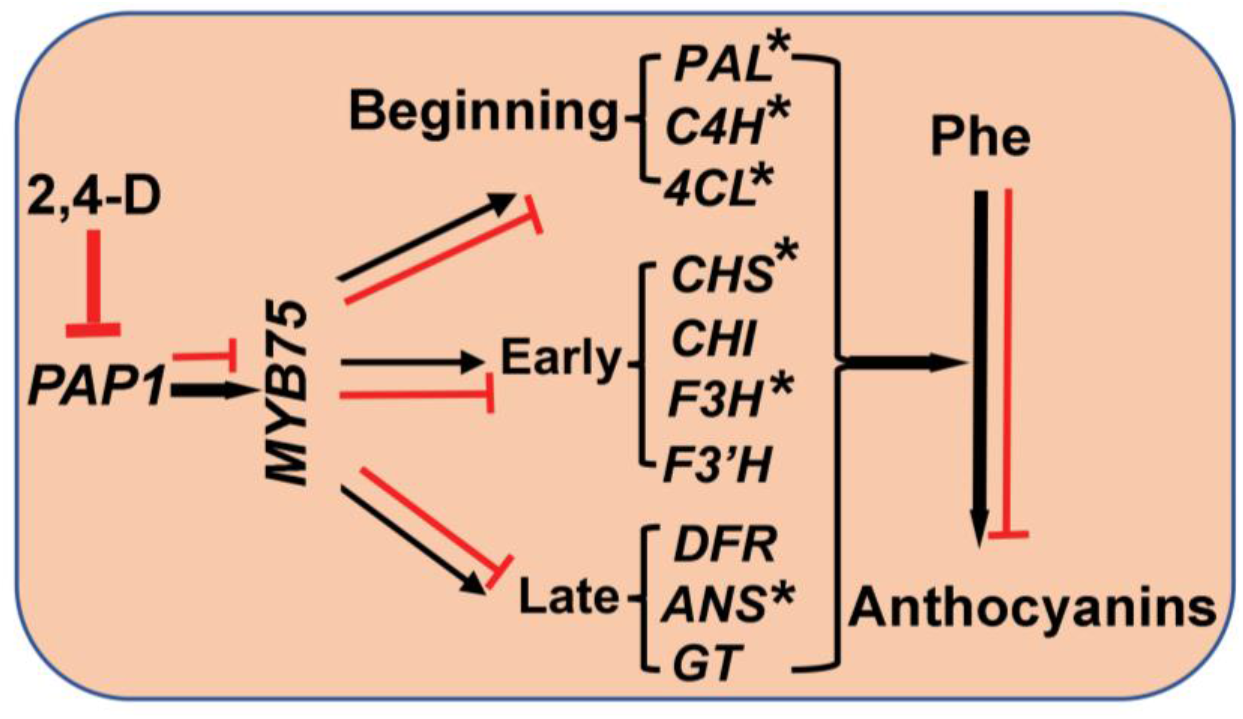
A simplified scheme characterizing the negative regulation of the biosynthesis of anthocyanins in TAPA1 cells by 2,4-D. Black arrows show the activation of the anthocyanin biosynthesis by PAP1. “Red T-shape lines or bars” means that 2,4-D downregulates the expression of the *PAP1* transgene leading to the reduction of the anthocyanin biosynthesis. “*” means that qRT-PCR data characterize their transcriptional reduction by 2,4-D.

## Abbreviation

KT: Kinetin
2,4-D: 2,4-Dichlorophenoxy acetic acid
NAA: 1-Naphtheleacetic acid
IAA: 3-Indoleacetic acid
MS medium: Murashige and Skoog medium (1962)
HPLC-TOF-MS/MS: High-performance liquid chromatography–time of flight–mass spectrometry
PAP1: Production of anthocyanin pigment 1
4CL: 4-coumaroyl: CoA-ligase
ANS: anthocyanidin synthase
C4H: cinnamate 4-hydroxylase
CHI: chalcone isomerase
CHS: chalcone synthase
DFR: dihydroflavonol reductase
F3H: flavanone 3-hydroxylase
F3’H: flavonoid 3’-hydroxylase

## Acknowledgement

We thank China Scholar Council for supporting Yilun Dong’s PhD Exchange Scholarship and Yue Zhu PhD study.

## Author Contribution statement

YD participated in the experimental designs of plant tissue and cell culture, performed all plant tissue culture experiments, extracted anthocyanins, performed HPLC-MS/MS analysis of anthocyanins and gene expression, and drafted material and methods, results, and discussion for manuscript preparation. ML mined genome sequences, identified promoters, analyzed cis-elements, cloned promoters and performed dual luciferase assays, and prepared materials, methods and results for manuscript preparation. BC and EY participated plant tissue culture, data collection and analysis, and anthocyanin extraction and analysis. YZ performed LC-MS/MS experiments and data analysis. LL and ZX participated in discussion of experimental design and data analysis. DYX perceived the entire project, designed and supervised all experiments, participated in data analysis, and drafted and finalized the manuscript.

## Data availability

All data that support our findings reported in this study are available in the supplementary materials of this article.

## Supplementary Materials

Sequences of eight pathway gene promoters and localization of AC and G-Box cis-elements.

