## Supplementary materials for "Molecular understanding of anthocyanin biosynthesis activated by PAP1 in engineered red *Artemisia annua* cells and regulation of 2, 4-dichlorophenoxyacetic acid"

**Nucleotide sequences of eight anthocyanin pathway gene promoters, AC-elements, and G-Box elements.**

G-Box

CDS-starting site.

AC-elements

*>AaPAL-pro*

AATCCAAATACCGTAAGGTGTATCTTGTTATAACCATCAATTCCACCTTCATGACGCATAATCGATAATGCTACTTATTTACCTAACTTTGCAATTTCGAAAGATTGTTTGTGAAATTACAATCTTATAGCTAAAAGTCTTTTGGTTCACAGGATTCTTAAAATTTTAATGAAATCCAAATAAATTTGTGGTGTATTCCTTTGTAGTGAGTAGGGCCTCTTTCTCGTTGACCAGTTCGTCTCCGCTCTATTGAAAGGTCTCCATTCCTACGGCTGTTTTGTCAATGAACCCGTGGATAAGTAGGCCTTTCTTGCAGACCGATCTTCTGCCGAGTTCCTTCCTTCGTAGTCAAATCCGATGATACGCTTTGGTGATGTTACCTATCAAATAAAAGGCCTACTAGGTGATCGAAATCGCTCAGGTCAGCGAGGACTCACTCACCTACTCCTTATCTAATACTTTCTTCTATCTACTAAATTAAAAAGCCGCCGTGCCGGGCTATAAGATAAGATACATCTGTTGTACTACTTGACTGGTAAGAAAGAGCTTCCGTAAGAACAGAAGACTCTTACATGTAGTAGGAATGCTACACTCACAAAAAAAATCCAAAATAAGGTGTCAGAAAGTGATTGGTGTATTTTGCTATAGTTTTTTTCTCTTCTTTTTGATTAAGTCGATCACTAATTCTCATTTTGTCCACAATTTTGTGAATTTTTTTTATGAGTGTAGCATCCTTTTATTTGAGTACAAAATCCGAATTACATTTCCATAGGTGATTGGTCGATAATTGAAAAAAAAAAAAAAAAAAAGAAAAAAAAAAAGAAGAAGCAAAATAGAGTTAAATTTGAAAGCAAGTTTTCAAGAGACAAGTTAAAATACTTGTACGAAGGAAAACAATAGTTTGTTTACTTCCCTAATTACAATATTTCTAAAACCCCCTAAAAAAAAGAGAAAAAAAATGCAACTCACCTACTACCACTAGCAATTTACGATTAGTGCATTGAATGAAAAACACCACTAGTTATTCAAGGTCCCGTATACCCACGTCATCGTAAATCCACACCATTAATTAAAACATACAAAATCCAACGGTCAATATTAAATCATGCAACCAACCTACCACCTTCGATCTCTTTTTCCACCCACCTACCACACTTCTCGTTATATAAACTACCACCTTACCACCCAATAAACCAACAATCTCAAAAAACATTATTATTTCACACACAAAGAGTGTGAGTCTAGTGTGTGAACTAATAACAAGAACAAAGATTGTTCTTTGTCTTATAATTGTGTTCTTTGTTTGCTCATTGTCTTTTGCACCAAATAAATAAATAAATAAACAACAATGGAAAACGGTACCGGGGTCTATGCTAATGGTAATGGTAATGGTCACATTAATGATTTATGCATTAAGGA

*>AaC4H-pro*

ACCAAACACTGGAATTTATATTAAATTTATGTGAGAATTCATTTCTGTGATTTGATTTTTGCACTTCATTTTATTTCTCTGAATTTCACCATATCAAACGTGCCTTCAAGAATTTAAAATTATAATTGAGAAATTCTAAAGAAATAGATTATGATTTAGTGAAGGTATGAATTATCTCTTATTTCATTTCAAATATTTTTTTTCAAATGTCACAATTAGTACAAGCTTAAAAATGAGTAGAATTTTTTTTTAGTTATCTATATCTATACTACATATAATACACAAATGAATCTCTCTCCAAAAATTTCAAATGTACAAAAAATTATGCAATGTTTACAACACTTTAAAGCAAGGAATCTTCATAATTACAACAAGTAAAGCATTGTGTACAAGCAAAACATGCATGACACGAAATTAAGACCAAAATACATAAATACATACATTAAATAGCCTTTAAAACACCCCTAAACCTAAAGTCACAATGATTTAAAGCATTGTGTACAAGCAAAACATGCACAACACGAAATTCAGACCGAAATACAGAAATATATACATTAAATAGCCTTTAAAACACCCTTACACCTAAATTTACAATGATTTAAACCTAAAGTACAACCCTCTATGAACATCTATTAGAAGCTTGACTTCGAAAACACAAAAGAAAACCTCATCATCGATCTATTAGAAGCTTGACTTATTTAAAGCGCTTATGGGGTGTTTGGGATTGCAGTGTTTGGGAAAGAAAATAAAAGGCTTGATTATTTGCGTTTCAGAAGCATCAAAACACGGTTTTGGAAAAGCAGATCCATACATGCTTTTTCAGAATGTGATTTTCCTTTCTTTGCCAAGTTACCGAACTGCCCGCATCCATCTTCTTCTTTGTTCATATTAATTTAACTTTTTGCATTTCTTATTTACCTATCTGAATATTTTGTTAGTGTGAGAAATAGATATTTTGAGAAAGACGAAAAGAGGCGGAGGCAGAACAAAAAGATGTAACCCTGATTTTAACTTCATATACTTCGTTATATCAGTATAAGCCCAGTAACAATAGATANTTGAAATCAAATTTGTTTAATTCACCGTCATACTTGACAAACTTCGTCACCAACTCCACCTCACCTAACCCCAACCTCTATAAAACAAACCACATTTCCAAAACACAAAACCAAAGAAAAAAAAAACACAAACACAAAAACACAACAACCATGGATCTCCTCTTCATAGA

*>Aa4CL-pro*

CTCCGGTGAGTAAGTCGTGTGGGGGTAGGTAGAGAAAAAGGCAAGCCAACCTGGATGTCAAACTTTAAAACATCACCCACCCAACTCAACCCAATAGTTGACCAACTCCAAAATGGCTCCTCAATATTAGAGTATTAAATCTAAAATAATAATATTACTCCCAATCTTACCTAACTCACACAACGTGACTTTAACTTTGGTAATCATGTTATATACTTGTATTAGCTTTATTTCTTTATTAAACACTTCACGTTATCTATTAACATGGTAAGACAATTAAATTGTGAATGTCACTCAAAGCTTTAATTTAAAGGATAAACATAACTATTCACAATTTTTTCTTTTTCCTTTTCTATTCCCTCATATTCATTGTATTTTAAACACATTTATACATTCATTTTTTTTCCAAGCCTATCAAAATATTTGAACTCAAGACTTTAATATCGAAAGGTTTAATATGAATCTCATCTCTATCGACGACTTATAAAATGTAGTTTTCATTGCGTTTAGACTCGTAGTATCTAATATGGGCTCGCAACCTTAGTATCGAAAGTTCTAATGCAAAGTCTGTCTTTATTGATGACTTGTAACATGTCTACCGTTCTAGTTTCTATCTGTCTCTATTAAAACCAATATTGAAAGAGAAGTGATATACATACACCAACTATTACGAGTAATTAAGATATACCACTATATAAACACCAACAAACATTATGTGACATATGTACAATTGTACTAACATTAAATTGTGTGTACAAATAAAAAAACAAACATTTACAAATCACCAACCATATCAAATGTTCTAAATATCAAAGTTCTAACATGAATTTCGTCTTTATTGATGACGTATAATACATCTAATCTTCTAGTTTTCTTCGAGTCGTAGAATCTAATATAAACTCAAGACTTTAAGGCAAGAAAAAAAAGAAAAGAAAATTGGACCCAATCACAAGCTCACCAACCACACCAGTGCCATAACATGAATTTCCATATCTACCCTTCAAATTAATACATAAAAATTCCCAATTTACCCTTAAATCACCAACCTTAAACTATATACCACGCGTATAAAGACCTTCACATTTTTATTACAACTTTTATTTATTTCCTATCAAACTCACCCACTTTCTTACCAAAGAAAAAAAATAACATTTCATTTTTCCGATGGCCTCACAACCGGAGCAAGAAAAAGAGATCATTTTCCGATCAAAGCTACCCGATATCTACATCCCAAAACATCTTCCTC

*>AaCHS-pro*

CTTACTTTCCTTTTCCCTTTCCAATTCTTTTCCATTTTCCTTCATTTAAAACTCGAGACCTGAGCGTTAGATATGTGCAAAAATAATCCTTATTTTTCCTATTTCCAGAAACATTTTCTTAGATATGTGCAAAAATTATTCTTATTTTGCTATTTCTAAAAATAATAATAATAATAAAAATAAATAAAATAAAAATCTAATAACAACGACACATACTCTGTCTTAAGAATGTTATATGTCAAGTATATTACTACAAGAGACAAACATGATAATAACAAGAAAAAGATGAAGTGTAAAAAATAGTACTGTAAACCGTTTGATAGTTTAGACCCGAAGAACACAATCAAGGAAAATGTTAAGGTCTGCGTATCTCACAGCTTTCTTAAGACAAATTCGTCCTCTCTATTACGAATGAATACACACATTAAAGCAAGATAAACAAAGGGGAGATCATATGATTATATTTCTTGAGAATATAACTCTCACTTTATAGTGATAGGAGATTAAGAAAAAGCAGTCGTATAATTAATCATACTAATGACTCCTTAACATAAAATTAATCCAGTCATTACATTTGACCATACATTAACTTCCATGTTAATAACTACTAGTATTATTGTAATTGATGACTATTAATTGTCATTCAAACCCATCATTTGAAAAAATTTCCTAACATTTCAAAGTCATTTAATGGCCCCTCCATAAAGTTTAAACTTTAAATTCCTAACAGTTCAAAGTCATTTAATGGCCTCCATAAGGTTTAAACTTTAAATGGCACTCTATCGAATATTACTTATGCACCTATAGTAAACTATATGTACCTTTTCTTATACATATCTACTTATGTTTATCCTAGCTTCAAAATCAAAAAATACTTGCAGAAAGTGATTGGTAAATATCAAGAAAAAATGGTCATTCACTTGATCTTTCCAATTTATTATGCATCATGAAACTAGTAAGTTGAAATCAGGTCTCCCACAAAAAGAGGGAAGATATCTGGTTTAAGATAACAAACAAATATGAATTTACTTATCCCATCACACTTCCAACACTTCAATGAACCCAAAACAAACCAAAGAAGTATTTATTAAGCTATTTAACAACAATTTCATGCTATTTCTTCTCTATACCAAACTTTCTCCAAGAACTGCCTATTTAATGCCTTATATATACACCCAAAACACCATTAACTTCTCACCACCAAACTTTAATCTCCTCAAGGTGAAGCAGCAGCATTCCTCACAAAATTATCCATGTCGACAACTAGGCGTCTTCCGACTCCAG

*>AaCHI-pro*

AATCAACACCTTCACCTTATTTTGTGTGTTGTGTGTTGTTCTTTCGTGTGTTTTTATTCGGTCATCAATTTACCAAATCAAATGAAAAGTCAGTGTGCCAATGGTGTAACACAAAAATGAACAAAACACGTGTGTTTTCATTCGCTCATCACTTTACCAAATCAAATGAAAAGTCACAAAGTGTCAATGATGTAACACAAAAATGAACAAAATGCGTGGTTATGATTCTTATTATGGTCAATCTAATCATCAAATATTTAACGATATAACACAAAATCAAAAGTTAAAAAACCAAGACAAATACCAAATTTTATAACCATTATTACTTATGGTGTCTTAAAATTTTCAAGGCCTTTACGTGAAATCTCACTTTAGGCCCCTCAGATCGTTAAGATGACCCAGTTTATTAAGGGAAATGTGCATCAAACATTCATATGCATGTTTTGAACATGACTACGCGATATTACATATATTTATACACGGTTATCATTGTGGTGCTACGTGCTAGACTCAAACCTAGATGGCGAAAAAGCTTTTTACATGTGTTTCATATAGATATTTCATGATAGACCTAAATCTATATAATCCAAAATTTCAATCATGGTGTGATAGTAATCTCTCTTCCCGAAATTCGAACTTATTATTAAAATAGTGTTATTTTATTCTCTTATACCATTGTGCTCACGGGCTTTGCGATTGATTTCCTGACATAAAAGTTCACTAAACATGTAGTCTTAACATAATCATATCTTAATCCTTACCAAAAAAATAAAATAAATTGGGTTCATCAGTATTGCCCCATATATGCAACATGTTTTTCGATTGTTGTTCCTATTAATGTAATTCAACTATCTTACTTTTTGTTTATTGATGAAGACACACAAGTCTTGAGGTTCATTTGTTTTGATTACTTTGTGAACGTTTACTATGCAGTTTAAGTAGTAAATCGTGTTGAGTTCGACAAAGATATGCATAAATCACGTTATTCTTATAGGAATCCTTTATTGTATATTGAATCTAAAACATAGTTTTACGTTTTTAATTAGCTATTGAGAAACTCGGCCATTGGCCGGGTAGAGACCTAGCATCAGGTACATCATATAAAATTTTGTTGAAGATATTAAATAGCAACACGTGTCATCATGGCAGTTTTAGGCTCATATTTAGGTTTGTTAATTAGGCTTGTTTATCATTTTCTTTTTAAAAGTTAGCATCAAGTTTGTCACATCGAGTCACTCGCTCATTATTTATCTCTCATTTACCCGATATTTATGCTCCTTTGAAAAAGAGAGAGTTTTCATTTCTGCTCCTTTTATCGTTATTGATACTTGATAGGAGTAGCATAATAATTTTTCAAACGTTCAATCTTCACCGTTAATTTCATCTGCCCCCTTAATTTTCTCCACACGATCTATCTACCACTTCAATTCCCTCCATAAATAATACTATTGATCAACATCTGCCAGTCAACATACAACACTTTTTCTTCACTCACCATGGCAACACCACCTTCAGCTAC

*>AaF3H-pro*

TTTTTACAAAGGGGTTGAGTATTTCTCAGCATAAACTGTTTTGTGATAAACTTGGGTTGGTGGATATGTTCAAGCCATGAGCTTGAGAGGGGGTGTTGAAATATATGGTTGGTCAAGCTCATGCATTGACTATTATAAGTTGAGGGACTTATTGTGTCAATATTATTCTCACTTTGTATGTTTAGTCTTGTATAGCAGGTTTGTTATATTTCAGGTACTGAAGTGTAACTTGTTTCTATATAAGCTGGGCTAACCCTAATTGTTCCATACAACACATTCAAGCCGTCCTATTTTTCACATACATCTTCTCTCATTGTGGTGGTTATCTCATTTTGTCTGGTCTTATAGGTTTGATCATCTTGATCAATAGTCTAGTATTTTCTGGTAGTATTTGAGAGTTTGTAGTGAGGCTGATTGTATTTGATTCTTTGTAGTGTGATAGATTAGACCCTTCTGAAGGGGACGATCTTTTCTCTTTGTATTTGTCTGGCAAGCACACGTTTTGTGGACTTGTCTTCTTGAACTATTAAGTAATAATAAGATTCACCAAGGGGTGGTTGGCTGGGTGATTTGTTAGTGTTGTAGTTGTTTTGAGATTAGATCTGTTATTTTAACAGTTTGACACGGTTTTGAAGCATTCTTCATAAATAAAGGTCGACAATATACGTATATAGAGTTATAGATAGTTTTCAAATCGTGTTTTTTTCGTCTTTGCTTTTTGGAACGTCCAAAAAAACTGCGTATTAAAAAACTAGTAACAAGCTTATGTTTGTTCTTAAGTAATGGAGTTTTATTATCCAATCATGTTACAAAGAATCACGTACGGCAACAAGTATAAGATATCGAATTCGATGATATATGATCGAAAGCTTTTGCAACGACATTCCTTACCTTCGATCACAATACATATACATAACGTAATCTTAGCGCAAACATTAACATGAACTCATTACATATTAGTACTTATAAACACTAAATGCAACATCTATTCAAGATTTTTTATAAGTAGACTCGATAAAAATCTTCGATAAGTGATGCAAAAACGTTATTGGGGTCACGTCTTGTAGGACCTTAGATCTAACTAAATCACTAATCTTTGAAGCACTTTGGTAACTTGAATGAAGTATATGTTCGACAAGTGAACCTCTGCCATGCTATTGTATGGCATAAACTTCTAGATCGTCAACCTTAAAAATATGGGGTTGATATCTAAAGCATTGATAATTACTTATATTTCTTGACTTTGCCTTTCCAAAAAAAAAAAACTTATATTTCTTGACTTTAAAAATTGCCATTTTAAAGACGGTACGCACAAAGTCTTTTTCTAGATCTTCAACCTTACAACTTTTATCTAGATATGTTAACTTTTCTTTCTTCTTTTTATTTTTGTGATAATTGTTAGAAAACTAAACATGGGTGTCCCACAACACTTTCTAT

>AaDFR-pro

TTTCTTTTTTGACCGTTTGGAGCTCTTTTAGAGTTGACCAAAAGGTTGACCTGGGAATAAAGTCTTTGTTCCTTCCTTCGAGTGGTTCTTGTGTTTGGTTTTTATTGAAGAAGAAGACTGTCTGCTATAGTCTTTTTCTACTTTGGAATTTTACGCAACTGTCTGTCTGTCCCTAAGATTTCGGGTATTGACGGATTAGTCCTTTGGTTTGTTTCTGCTTTGTAACATGTTCTAATAGTTTGTAACATGTTCTAGTTTACTTGACTCTATAATGGAATTTTAGTTTTTTTTAATTTTAATGGAATTAAAAAAAAAAAAGTGAGTGCAGGGTGAGTGATCGGATGCCAACAAAAATATGGGTGAGTGATGAAAAAAGTGGGTGAATGATGTGGCATAAGGAGATTGGTTGGGAGTGAGTGATGGGAGAGAGAGTGGGTGATCCACTGCCAACCTCCTTAGCTTATTGAAGTCTACAATATATGATAGATTGTCGATCAAAAAAATCTTATATTTAATTACTTGTTTGCAATTTCATACGGAGTAACATTTAACTACTTCAACGATACTAAGCAATACTAGAATGCACATTTTTTTTTAAACATGAATACACTAACTTTTGAACCAAGTGTAGTAAAAACTTGTTTTCTACGTCCAAACTAGCAATGCGCGACAAAAAAACATATATGTCTCAAAAGGCACTTTCACTCCAGCCTGATACAAGTTAATTTGCACATGTACAAGTACAAGTATGACCACCACTAAAAAACCGTACTAATGACCACGATCATCATCACCACGTGTGATGATGTGATTTTGGTGTCGACTTGGTAGCTGAGGCGACCGAGTGGAGAAAGACAAAAATGTGCGAGTTTGAGAGTGTATTTGGTGGTGTTAGTTTGGATTTAGTGTTTAATTGGATAGACATTACTATGACCCCTTAGTGATGGATTTGATAAAAGAGTAAAACACTGACGTGTCTAAGACTAAGTGTGATAACTCGGTCGACTCTCGGTCGGTAGAGGGTGCTACGCGTCGGCCCTTAATTACTAGGGCGGCATATAGGAACGAAATGGAGATGTTGGTCGCAAAAGCGACTCACTTTTGTGCTTGTCTGTATGTGTGGGAGTTTAATTTAAAATATATTTTTGTAGGGTATGTTTTTGTGTTATCGTTCTCCTTATTACACTAAGTTGATAATGAAATAGTCTTGAAGATAAAGATAGAAAGTTAATGTGTTGTGGTTGTCATTACTACACCCGGTTTAAAAAGGCAGTAAAAGTTTGGAATTTTATGGCGAATTTCTGTGAAATAGTCTCAAACCCAACGGTCAAACGTTGCATGCAACCATCCTCACCCCCACCATTCGATCGAGTTGAATGGTGGCTTCTGCATGTGCATTAATTTGTCTAATTTTTTATACATTAATACGTATACTAAAAAACAAATACTTGAGAAATGACAAGAGAATTGAACTCTTTCTATTCGGGAGTTGAATGGTGAGTTCAGCACGCGCGTTTATTAGATACTCCGTATATATTAAATCCTTCTATAAATACTCATCTCGATTATTACTTAATCCCATCATCATCCACAAATACTTCAAATACCACAAGATTAAAACAACATGAAAGAAGACTCACCA

>AaANS-pro

GGGTGGTTAGGTTATCCGAAATGATACGATTATTCGGATGATGATCTGGGGTGTATTGGTTTGATCTGTCCGGACATCAGTCTGTTCCGTCCGGATGTTAGTCTTTCGCGTATCATAATTCAGAGGTTTTATTCCGGGGTCTCGTTCCGGGGCTTCGTTCCTTCCGGACGGGGTCCGGAGGACACATAAGCCAACAATCTGCATTTAAAAAGATGAAGCGAGAGTGTTGGTAATCGAAGTAAGATTTCCATCAAAAGGTCGTCATTGGATAACACTTTATATACGGATGAAGGGAAGTTGAGGTTTATGCCATCATCATCATTGGCAACAACCTCCAACCGGAGGGGAAGTCCAGATGAACCACCGGCCATCGTTTTGTAAATCAAAACTCTAACGTTTGGGTTCTCCGAATTTGAGTTTTAGGGTTTTGATTTGGGGACCTTTATTTAAAGGCTAAAAAAATATGCATGTTTTAAAAATTTAAGTTTCGATCAACTCACGATTTTAACCCTCCAACTTTCTTGATCATTGATGTAATTCTTAACTTTTGTTGGCTTGTTGAAATAGCCATCAAAACATACTATTGAACAATAATAGTTTAAATTTAAAAATGATTTTAGATTTCTGTGCATGCACGGTGTTTTTGAGATAAATTTTTGCTTAGAGTGTTTATACACGTTCGATAAATTTACTTGTATATTATTAAACACAAGGTAAAGGATAATTTAAGAAAATTTTGCTACATTGCAACTGGGTAATTTTTTGTTGGATTTCACTTCTATAAATGTCACATAGTGAAGCTGAGGAACACGAAACACCTAATACGAGGTCGTCGCATAATTATGTTAGACTTTGGAATGGTCGATTTAGTTGGATGGTTACTGGACTTTAATGGTTCATCAAAAAATGAGGCGATTAGTAAATATATAAAGTAAAAAATAAAAAAACTGAATGATGTTTAATTTGTCATAAACAAACAGGTATCCTACTTCAATTCCTTATGAATATCCACAAAATTGTTAAAATTCATGAAAGGATGAATCATTTGTTACGACAAATTAAAAAGGTGTTAAATTTTGGATGTTGGTTGGTAAACTTGTTCCCACTCGAACGAAGAGAACAATATGTGTATATATACTCCTTTAGTAGTACCCCTAATACAGATCTATTGGTTTTTATTATCGAGTAAAAATATATTAGCTATGTTACTGATAATTATTAAAAAATCTTTTACTCAAAACTTGTTTTTTGACAATTTGTTTTTATTGGTCACTCTTAACTTGTTTTTACTTATTAAGAGTCAAATTTGACTTAATTCTAATACGTTGAAACCATCTTTTTATTGACGGCCAACATAAAATGTTTCAGTAATCAAACTTTAACATGAAATTGTTTGTGTTGAATTTGAACACCACGTTTCTTAAGCTGACCTTTCAAGANAACCTACCATTCTTAAAAACTTCAATATATACATATAAATGTACTCTCCATATAATCACTAAAAGAAACACCACAAAACCTTCAAAATACAACACTTACATACAAAATGGTGATTCCAACAAACACAAGA

Table S1 Primer pairs designed for qRT-PCR and thermal cycle programs

| **Gene** | **Primers for RT-PCR and/or qRT-PCR** | **Thermal Cycle Program for RT-PCR** |
| --- | --- | --- |
| *AtPAP1* | Forward: 5’-GACATTACGCCCAT  TCCTACAAC-3’  Reverse: 5’-TCGAGGTCGAGGC  TTATAAACATT-3’ | 95°Cx2’ + 35x(95°Cx45”+ 55°Cx 45” + 72°Cx1’) + 72°Cx5’ |
| *AaPAL1* | Forward: 5’- CTCTTCCACATTCA  GCAACACGA-3’  Reverse: 5’- TTTTCAGCAGTCAGG  ATTTCACC-3’ | 95°Cx2’ + 25x(95°Cx45”+ 55°Cx 45” + 72°Cx1’) + 72°Cx5’ |
| *AaCHS* | Forward: 5'-CGATTACCAACTCAC  AAAACTC-3'  Reverse: 5'-GTCAATATGGTTT  TCATTAGGC-3' | 95°Cx2’ + 35x(95°Cx45”+ 55°Cx 45” + 72°Cx1’) + 72°Cx5’ |
| *AaDFR* | Forward: 5’-GGAGTGTTTCATGTT  GCCACCCCTATGGA-3’  Reverse: 5’-CCGGAATTTGC  TGTTTTTGTGCATTAACAGTCCC-3’ | 95°Cx2’ + 30x(95°Cx45”+ 55°Cx 45” + 72°Cx1’) + 72°Cx5’ |
| *AaANS* | Forward: 5’-TGTGTACCAGACTCC  ATCATCATGCACATTGG-3’  Reverse: 5’-CGGAAAGAGTGGTGG  GTTCTCCTTAGAAATGG-3’ | 95°Cx2’ + 30x(95°Cx45”+ 55°Cx 45” + 72°Cx1’) + 72°Cx5’ |
| *AaBeta-Actin* | Forward: 5’-CCAGGCTGTTCAGTCT  CTGTAT-3’  Reverse: 5’-CGCTCGGTAAGG  ATCTTCATCA-3’ | 95°Cx2’ + 35x(95°Cx45”+ 56°Cx 45” + 72°Cx1’) + 72°Cx5’ |

Table S2 primers for amplification of eight gene promoters and restriction enzyme sites added for cloning.

| Promoter primers | registration enzyme site | Sequence | Thermal cycle |
| --- | --- | --- | --- |
| Aa 4CL2 pro F | PstI | **AAACTGCAG**CTCCGGTGAGTAAGTCGTGT | 98°C x 30s    98°C x 10s  60°C x 30s, 30 cycles  72°Cx 45s    72°C x 5 mins  4°C |
| Aa 4CL2 pro R | Smal I | **TCCCCCGGG**GGTAAGAAAGTGGGTGAGTT |  |
| Aa PAL pro F | PstI | **AAACTGCAG**TAACCATCAATTCCACCTTC |  |
| xAa PAL pro R | Smal I | **CCCCCGGG**GCAAAAGACAATGAGCAAAC |  |
| Aa C4H pro F | Pst I | **AAACTGCAG**TTATTTCTCTGAATTTCACC |  |
| Aa C4H pro R | Smal I | **TCCCCCGGG**GGTTGTTGTGTTTTTGTGTT |  |
| Aa CHS proF | PstI | **AAACTGCAG**TACTGTAAACCGTTTGATAG |  |
| Aa CHS proR | Smal I | **TCCCCCGGG** GAAGACGCCTAGTTGTCGA |  |
| Aa CHI2 pro F | PstI | **AAACTGCAG**TTTTTATTCGGTCATCAATT |  |
| Aa CHI2 pro R | Smal I | **TCCCCCGGG**AAGTGTTGTATGTTGACTGGC |  |
| Aa F3H pro F | Pst I | **AAACTGCAG**GAGTATTTCTCAGCATAAAC |  |
| Aa F3H pro R | Smal I | **TCCCCCGGG**GTTTAGTTTTCTAACAATTA |  |
| Aa DFR pro F | Pst I | **AAACTGCAG**ACTACTTCAACGATACTAAG |  |
| Aa DFR pro R | Smal I | **TCCCCCGGG**TGTTTTAATCTTGTGGTATT |  |
| Aa ANS pro F | Pst I | **AAACTGCAG**ATTATCGCTAATCTCGGGTG |  |
| AaANS proR | Smal I | **TCCCCCGGG**TTGTATTTTGAAGGTTTTGT |  |
